# Niche separation in cross-feeding sustains bacterial strain diversity across nutrient environments and may increase chances for survival in nutrient-limited leaf apoplasts

**DOI:** 10.1101/2021.11.07.467568

**Authors:** Mariana Murillo-Roos, Hafiz Syed M. Abdullah, Mossaab Debbar, Nico Ueberschaar, Matthew T. Agler

## Abstract

The leaf microbiome plays a crucial role in plant’s health and resilience to stress. Like in other hosts, successful colonization is dependent on multiple factors, among them, resource accessibility. The apoplast is an important site of plant-microbe interactions where nutrients are tightly regulated. While leaf pathogens have evolved elaborate strategies to obtain nutrients there, it is not yet clear how commensals survive without most of these adaptations. Resource limitation can promote metabolic interactions, which in turn shape and stabilize microbiomes but this has not been addressed in detail in leaves. Here, we investigated whether and how the nutrient environment might influence metabolic exchange and assembly of bacterial communities in *Flaveria trinervia* and *F. robusta* leaves. We enriched bacteria from both plant species *in-vitro* in minimal media with sucrose as a carbon source, and with or without amino acids. After enrichment, we studied the genetic and metabolic diversity within the communities. Enriched *Pseudomonas koreensis* strains could cross-feed from diverse leaf bacteria. Although P. koreensis could not utilize sucrose, cross-feeding diverse metabolites from *Pantoea* sp ensured their survival in the sucrose-only enrichments. The *Pseudomonas* strains had high genetic similarity (∼99.8% ANI) but still displayed clear niche partitioning, enabling them to simultaneously cross-feed from *Pantoea*. Interestingly, cross-feeders were only enriched from *F. robusta* and not from *F. trinervia*. Untargeted metabolomics analysis of the leaf apoplasts revealed contrasting nutrient environments, with greater concentrations of high-cost amino acids in *F. trinervia*. Additionally, *P. koreensis* strains were better able to survive without a cross-feeding partner in these richer apoplasts. Thus, cross feeding might arise as an adaptation to cope with nutrient limitations in the apoplast. Understanding how apoplast resources influence metabolic interactions could therefore provide plant breeders targets to manipulate leaf microbiome shape and stability.

## Introduction

In nature, plants are colonized by diverse communities of microorganisms. A balanced microbial community can work as a barrier against biotic and abiotic stress, helping to sustain plant health. On the other hand, failing to establish a normal microbiota can have devasting effects (1). In-depth descriptive studies have shown that the composition of the plant microbiome depend on factors like host species/genotype, tissue type and development stages, and the abiotic environment (2–4). A significant part of these differences can be attributed to networks of interactions among microorganisms (5,6). Besides shaping the microbiome, these interactions play critical roles in maintaining their stability, for example to invaders in both roots and leaves (6,7). Plant-associated bacteria are highly diverse, whereby single taxa can have many genetically distinct strains within a single plant population (8). While this diversity probably arises in part because of interactions with plants, the role of the extensive microbe-microbe interactions all colonizers face is still unclear. Given the importance of these interactions, increasing the understanding of how they arise and how diversity influences them, may help develop better strategies to increase plant resilience.

Microbial diversity in plant tissues generally follows a gradient from high in roots to lower diversity in leaves, with relatively few taxa that further colonize the leaf apoplastic space as endophytes (1,9,10). This limitation is due to strong constraints on microbial life in the apoplast, including the tight regulation of nutrient resources imposed by plants, presumably in part to limit microbial growth (11–13). Bacterial survival requires at least basic nutrients like carbon and nitrogen. Both sucrose, the primary sugar produced in leaves, and amino acids are found in the apoplast (14,15) and can play important roles in bacterial nutrition and virulence (16–18). However, their availability is unstable and varies both between plant species and within plant species, for example due to diurnal fluctuations (19).

The lack of resources in plant apoplasts is evident when it is considered that, an apparently fundamental trait of bacterial leaf pathogens, is the ability to mobilize resources using for example batteries of secreted effector proteins (20,21). Non-pathogenic strains (i.e., commensals), however, have fewer adaptations to manipulate plant nutrient availability and therefore are probably more directly reliant on scavenging nutrients (22). For example, commensal *Burkholderia* and *Agrobacterium* express diverse transporters for ribose, xylose, arabinose and urea transporters upon injection into *Arabidopsis thaliana* leaves (12). For colonizers like these, metabolic interactions with co-colonizers are likely to play important roles helping them gain nutrition. Specifically, resource limitation increases the likelihood that cooperative or even mutualistic interactions arise, including metabolic cross-feeding (23–25). Such cooperative interactions can also be beneficial by making communities more resilient to fluctuating nutrient availability (26). Although it is speculated that microbe-microbe interactions in leaves involves nutrient exchange, it is not yet clear how widespread this is and whether cooperation may impact the establishment of the leaf community (27).

Here, we investigated how the apoplast nutrient environment may influence the arisal of metabolic interactions among bacteria. As a model system, we compared plant species in the genus *Flaveria* that use different photosynthesis strategies (C3: *F. robusta*, C4: *F. trinervia*). Although they are closely related, the evolution of C4 photosynthesis has had direct and indirect effects on traits like leaf metabolism (28,29) and leaf structure (30). Thus, we hypothesized that these differences would be likely to affect the arisal of metabolic interactions. To address this question, we used an in-vitro community enrichment approach and dissected inter-bacterial interactions at the strain level using metabolomics, genomics and molecular tools. Diverse leaf bacteria have the potential to cross-feed and we found that cross-feeding potential correlated with a more nutrient-poor apoplast environment where survival of individuals is limited. Additionally, we found that metabolic interactions sustain taxonomic diversity across vastly different nutrient regimes and that partitioning of cross-feeding niches can sustain strain-level genetic diversity. Thus, our results suggest that metabolic interactions help bacteria cope with nutrient limitation in host plants, and that leaf traits have the potential to shape leaf microbiomes.

## Methods

### Data and Resource Availability

Sequencing data is publicly available on NCBI SRA (Project number: X). Metabolomics data is publicly available on Metabolights (Project number: X). Scripts to process raw data, analyze data and generate figures is available on FigShare (Public website: X).

### Enrichment of leaf microbiomes from *Flaveria trinervia* and *Flaveria robusta*

Cuttings from *Flaveria robusta, F. linearis* and *F. trinervia* were grown in an outdoor garden for two months(Jena, Germany) to allow natural colonization by microorganisms. Well-developed leaves were sampled, weighed, and washed three times in sterile water to remove dirt and insects and leaf microbial extracts were prepared by macerating the washed leaves in 1X PBS + 0.02% Silwet and adding 20% glycerol before storing at -80 °C. About 1000 cells (estimated by plating) were pre-cultured in M9 broth with trace elements, 11 mM sucrose, 0.2% w/v casamino acids (Difco) 200 mM NH_4_Cl and 200 µg/mL of cycloheximide to limit eukaryotic growth. After washing cells and standardizing OD, the pre-culture was used to inoculate the two enrichment M9 media supplemented with NH_4_Cl (33 mM) and either no casamino acids (S-CA) or 0.2% m/v casamino acids (S+CA). For each 48-h passage of the enrichment, 5uL of homogenized enrichment was transferred to a new plate with fresh media. At the last passage, cells were collected for DNA extraction and to prepare glycerol stocks stored at −80 °C. Details of the entire procedure can be found in the Supplementary Methods.

### Characterization of bacteria in original leaf extracts and in enrichments

To isolate and identify bacteria in the communities, 25 isolates were recovered from the initial leaf extract glycerol stocks and from the glycerol stocks from the twelfth enrichment passage of each condition (150 total). All isolates were identified *via* Sanger sequencing of the 16S rRNA gene. Additionally, bacterial communities in the twelfth enrichment passage were characterized by 16S rRNA gene amplicon sequencing of the V3-V4 region similar to the two-step approach outlined in (31). Data was analyzed in R using the packages dada2, phyloseq and vegan.

Whole genome sequencing was performed on three *Pseudomonas koreensis* (*Pk*Fr_-CA_5,_ *Pk*Fr_+CA_3_ and *Pk*Fr_+CA_18_) and four *Pantoea sp*. isolates (*Pa*Fr_-CA_6_, *Pa*Fr_+CA_20,_ *Pa*Ft_-CA_14,_ and *Pa*Fr_+CA_17_). For this, DNA was extracted, purified and sent to Microbial Genome Sequencing Center (Pittsburgh, USA) for sequencing on the NextSeq 2000 platform at a depth of 300 MBp (∼50x for P. koreensis strains and ∼60x for P. agglomerans strains). The genomes were each assembled using SPADES (3.14.1) and average nucleotide identity (ANI) between the different isolates of each genera was calculated in Kbase (32). Single nucleotide polymorphisms (SNPs) compared to the *Pseudomonas koreensis* D26 reference genome (assembly accession GCF_001605965.1) were called by mapping the raw sequencing reads from each *P. koreensis* strain using SNIPPY (version 4.6.0). Full details on DNA extraction, PCR and data analysis can be found in the Supplementary Methods.

### Evaluation of cross-feeding interactions among isolates

After testing all isolates for their growth patterns in the media they were enriched in and for potential auxotrophies (see supplementary methods for full details of evaluation of isolate growth) we identified some strains must have cross-fed and this was investigated in-depth. To generate spent media, sucrose-utilizing bacterial isolates were grown in 250 mL flasks with 80 mL of S-CA (sucrose-only) media at 26 °C and 220 rpm. After 48 hours, the cells were separated from the spent media by centrifuging at 5000 x g for 5 minutes and filter sterilizing twice through a 0.22 µm PES filter. Spent media were subjected to untargeted metabolomics or used further to assay growth of dependent bacteria. Consumer bacterial strains were grown in R2 broth for 24 hours at 28 °C and 220 rpm; the cells were washed twice and resuspended in 1x PBS to an OD of 0.2. In 96-well plates, 180 µL of the sterile spent media and 20 µL of the bacterial suspension were mixed (final OD 0.02). As a negative control, the isolates were also inoculated in full strength S-CA media. After or during 48 hours of incubation at 28 °C and 220 rpm, growth was measured (OD_600nm_). To measure taken up compounds, the plate was centrifuged at 5000 x g for 5 min, and the supernatants were transferred to a clean plate and stored at -20 °C until injection in the HPLC-MS. To test for inhibitory effects, the spent media of the isolates *Pa* Ft_-CA_14_ and *Pa* Ft_+CA_17_ were diluted 1:2 with spent media of *Pa* Fr_-CA_6_ before growth of the consumer strains.

### Competition experiment using tagged *Pseudomonas* strains

To evaluate cross-feeding of fast and slow-growing *Pseudomonas* (*Pk*) isolates together with *Pantoea* (*Pa*), we first tagged the isolates *Pk* Fr_-CA_5_ and *Pk* Fr_+CA_3_ with the fluorophores mTagBFP2 and mOrange2, respectively, making them clearly distinguishable for colony counting. We used the delivery plasmid systems pMRE-Tn7-140 and pMRE-Tn7-144 developed by Schlechter et al. (33) (see Supplementary Methods for details). Tagged *Pk* strains and *Pa* Fr_-CA_6_ were precultured in R2 broth and diluted to an OD of 0.2. In black 96-well plates, 180 µL of S-CA media were inoculated with 20 µL of the corresponding bacterial suspension in quadruplicate (Supp Table 1) and covered with an AeraSeal™ (Excel Scientific, Inc.) sealing film. The fast-growing *Pk* isolate was added at half the density of the slow-growing isolate. The full community (both *Pk* strains together with *Pa*) was sub-cultured every 24 hours or every 48 hours for a total of eight and four passages, respectively, by transferring 5 µL to 195 µL of fresh media. The controls, i.e., each *Pk* strain alone with *Pa* and *Pa* in monoculture were only grown for the eight 24-h passages. The plate was incubated in a BioLector I (m2p-labs Beasweiler, Germany) at 500 rpm, 30 °C with humidity control. After each round, the plate was opened under sterile conditions, and 2 µL was sampled for serial dilutions and CFU counts, whereby *Pk* Fr_-CA_5_ and *Pk* Fr_+CA_3_ were distinguishable due to fluorescence. During the runs, we monitored growth of *Pk* Fr_+CA_3_ by normalizing the red filter channel signal (set at gain=100) to the biomass signal (set at gain= 35), so that we could compare its growth in co-culture with *Pa* alone vs. together with *Pk* Fr_-CA_5_.

### *In-planta* testing of *Pseudomonas* colonization

To test the ability of the fast- and slow cross-feeding strains (*Pk* Fr_-CA_5_ and *Pk* Fr_+CA_3_, respectively) to persist in leaves of *F. robusta* or *F. trinervia*, we inoculated three-weeks old cuttings grown in potting soil with a washed cell suspension of either *Pseudomonas* (*Pk*) alone or in combination with *Pantoea (Pa)* (OD 0.002 of each). The plants were arranged in trays, and these were covered with a plastic lid to keep a humid environment and placed inside a growth chamber with a photoperiod of 16 hours and a day/night temperature of 22/18 °C. The plants were harvested ten days after inoculation and leaves processed to count CFU/g of leaf. The inoculation was repeated in two fully independent experiments for each species. See Supplementary Methods for full details.

### Recovery and metabolomic analysis of leaf apoplast fluid from lab plants

Cuttings from *Flaveria robusta, F. linearis* and *F. trinervia* were grown in potting soil as described above. The plants were kept at an average day/night temperature of 25 °C/22 °C and a photoperiod of 16 hours. Apoplast fluid was extracted from fully developed leaves by infiltrating them with sodium phosphate (100 mM, pH 6.5) under vacuum in a syringe and recovering it by centrifugation, similar to Gentzel et al. (34) An infiltration ratio, used later to correct the metabolite peak areas for the dilution that occurred during infiltration, was calculated by dividing the mass of buffer that went into the leaf over the initial weight (*W*_*inf*_ -*W*_*ini*_)/ *W*_*ini*_. After storage at -20 °C, the samples were spiked with an internal standard of deuterated amino acids and subjected to metabolomic profiling via untargeted UHPLC-HRMS (Supp Table 9). Full details on the apoplast recovery as well as UHPLC-HRMS parameters and data analysis can be found in the Supplementary Methods.

## Results

### Nutrient and taxonomic diversity shapes microbiome function in *Flaveria* leaf enrichments

We first tested whether nutrient richness influences diversity and interactions in communities of leaf bacteria. We enriched bacteria derived from leaves of *F. trinervia* (*Ft)* and *F. robusta* (*Fr*) (Fig 1a) through sequential 48-h passages in a base M9 minimal media with sucrose and ammonia and either no amino acids (S-CA) or addition of casamino acids (S+CA) at low levels similar to those previously reported in the apoplast (35). The OD attained at the end of the experiment was used as a measure of productivity (function) of the enrichments (Fig 1b). In the S-CA condition where sucrose was the only carbon source, the *Fr* enrichment was slightly more productive than the *Ft* enrichment. Additionally, enrichments from *Fr* tended to increase their productivity when the potential carbon and nitrogen sources were richer (S+CA), but those from *Ft* did not.

**Figure 1.**
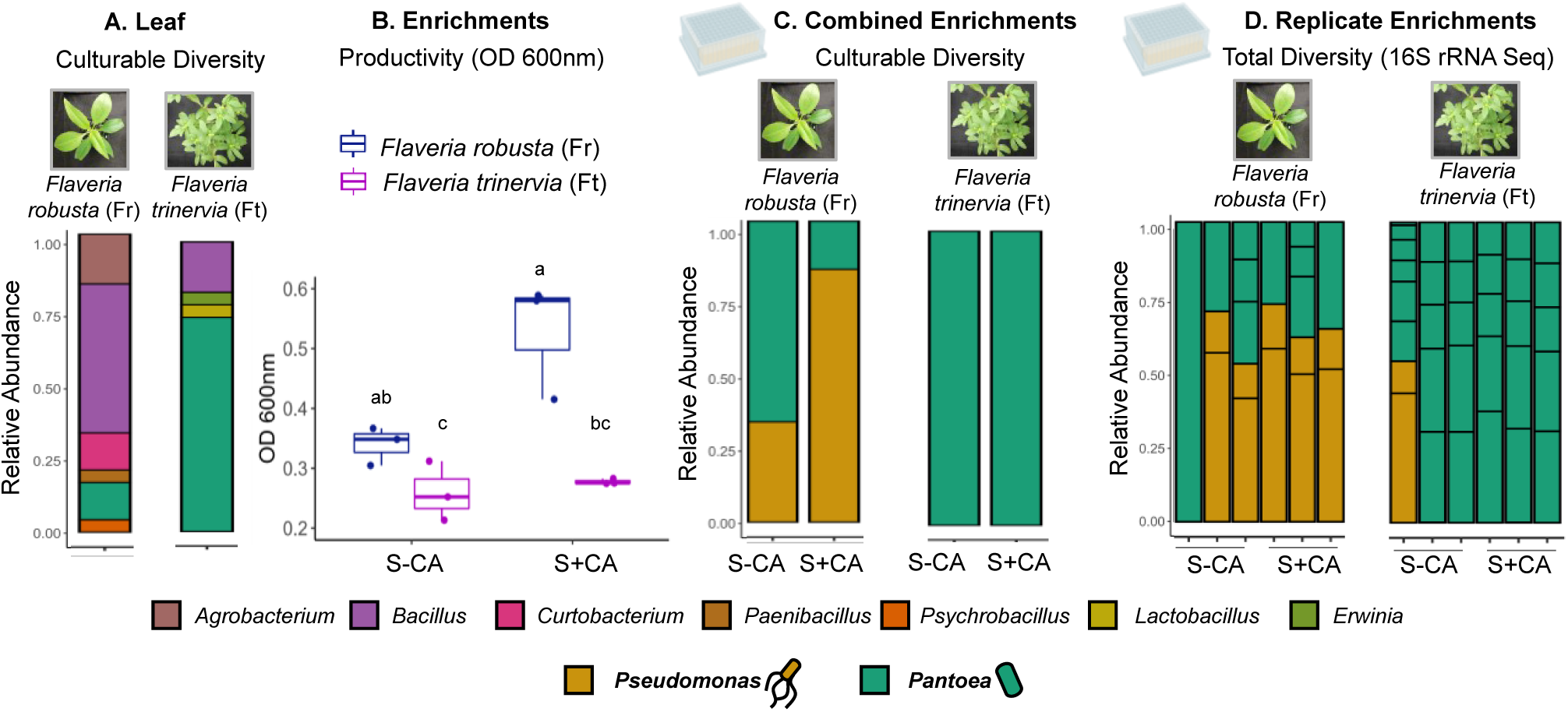
Bacterial community enrichments from *F. robusta* leaves show higher productivity and more functional responsiveness to nutrient diversity than *F. trinervia*. A) Culturable diversity in the leaf extracts used for enrichment from *Flaveria robusta* and *F. trinervia*, based on 16S rRNA Sanger sequencing; n=25 isolates. B) Productivity (final OD) of the enrichments from each species. Different letters indicate significantly different values (Kruskal Wallis test with FDR adjustment <0.05). (N=3) C) Culturable diversity obtained in each enrichment after 12 passages based on Sanger sequencing of 16S rRNA gene (Three replicates were combined for isolation); n=25 isolates. D) Diversity obtained in each enrichment replicate after 12 passages based on amplicon sequencing of 16S rRNA gene. The taxonomy color legend corresponds to panels A, C and D.

To investigate why the productivity differed between the enrichments, we collected 25 random bacterial isolates from the S-CA and S+CA communities, identified them taxonomically (Fig 1c) and tested the growth of each in S-CA and S+CA (Supp Table 2). Additionally, we conducted amplicon sequencing on the full communities to have a broader picture of their composition (Fig 1d). Surprisingly, despite diverse bacteria in the original leaves (Fig 1a), enrichments selected very few taxa, which was not affected by additional resource availability in the form of amino acids. *Ft* enrichments were fully dominated by *Pantoea sp*. (hereafter *Pa*) (Fig 1c and 1d). *Fr* enrichments also contained a high prevalence of *Pa*, but in combination with *Pseudomonas koreensis* (hereafter *Pk*) (Fig 1c and 1d); in the S-CA community, the fraction of *Pa* was higher than in S+CA (67% vs. 21%, respectively, Fig 1c). All tested *Pa* isolates could grow on sucrose or amino acids as a sole carbon source. The fact that productivity did not increase in *Ft* enrichments with casamino acid addition suggests that nitrogen may have limited additional growth. All tested *Pk* isolates could use amino acids as a sole carbon source. Thus, the increased productivity in the *Fr* S+CA enrichment suggests that they could make use of the additional resources. Surprisingly, *Pk* isolates did not use sucrose as a sole carbon source even though they were present in the S-CA (sucrose-only) enrichment (Supp Table 2). Therefore, *Pk* in this enrichment may have been somehow dependent on *Pa*, which could explain the higher productivity in *Fr* enrichments compared to *Ft* when sucrose was the only carbon source.

### Cross-feeding on diverse metabolites sustains *P. koreensis* in the absence of a primary carbon source

We asked how *Pseudomonas koreensis* (*Pk*) isolates survived in the S-CA enrichment if they could not utilize sucrose directly. One possibility is they were auxotrophs and could grow when amino acids were available in the environment (either in S+CA or provided from *Pantoea* (*Pa)* in S-CA). However, all tested *Pk* isolates could grow on glucose without amino acid supplementation (Supp Table 2), suggesting that they were not strict auxotrophs.

Next, we tested whether *Pa* produced other metabolic by-products that *Pk* may grow on. Indeed, all *Pk* isolates from the sucrose-only enrichment (*Pk* Fr_-CA_) could grow on spent media of a *Pa* isolate from the same enrichment (*Pa*Fr_−CA_6_, Fig 2a). To determine what *Pk* consumed, we performed untargeted metabolomics on the spent media before and after growth of three *Pk* isolates and considered metabolite peaks to be consumed if their area strongly reduced after *Pk* growth (log2FC<-2, FDR<0.05). With this strict cutoff we detected uptake of between 23-25 metabolites by each strain, including hypoxanthine, spermidine and, in one of the isolates, alanine. (Supp Table 3). With slightly looser parameters, (log2FC<-2, p.value <0.05) we also found uptake of N-acetylputrescine, guanine and maleic acid (Supp Table 3), indicating a complex metabolic dependency of *Pk* on *Pa* in the sucrose-only enrichments. We did detect trace levels of glucose and/or fructose in the *Pa* spent medium (hexoses are indistinguishable with this method), but they were not significantly taken up (log2FC=-0.85, p.value=0.36 in average, not shown).

**Figure 2.**
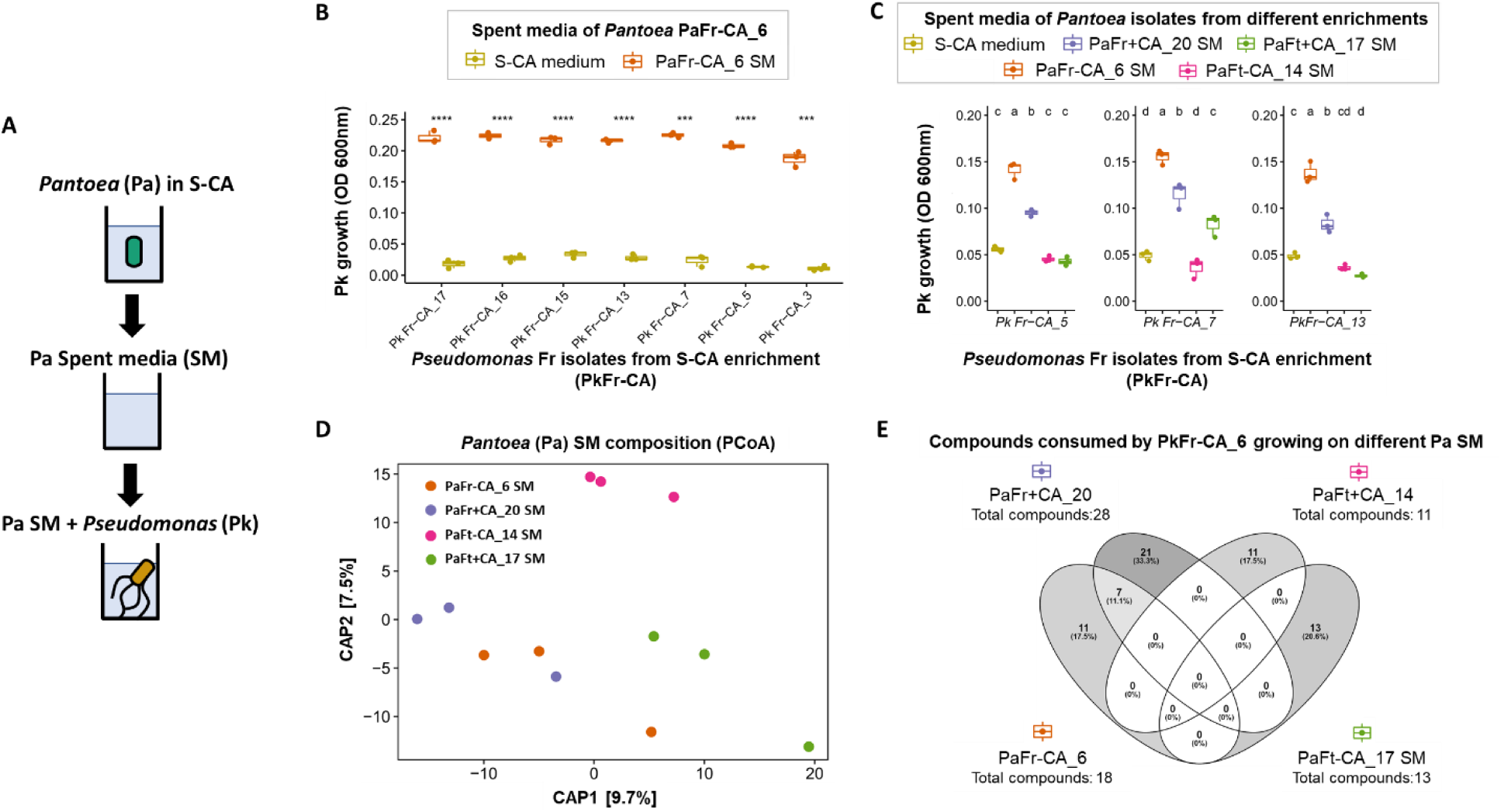
*Pa* Fr isolates were better cross feeders than *Pa* Ft, although their spent media were not extremely different. A) Setup of cross-feeding experiments. B) *Pk* Fr-CA isolates growth on *Pa* Fr-CA_6 spent medium. (N=3) C) *Pk* -CA isolates growth on different *Pa* isolates spent media. (N=3) D) Constrained analysis of principal components of all peaks (area>1.0E4) detected in the spent media of the different *Pa* isolates. The data was glog transformed and the analysis was based on Euclidean distance matrix. (N=3) D) Venn Diagram of significantly taken up metabolites by *Pk* Fr-CA_5 from the different *Pa* spent media (log2FC<-2, FDR<0.05). (N=3)

Interestingly, we observed that several isolates from the original leaf extracts used to inoculate the enrichments (Fr_LE_ and Ft_LE_) could also feed *Pk*. The growth on *Bacillus aryabhattai* Fr_LE_12_ spent medium, for example, was especially high, even compared to growth on *Pa* spent medium (Supp Fig 1c). Metabolomic analyses suggested that *Pk* may utilize a diverse array of metabolites from this isolate (log2FC<-2, FDR<0.05) with little or no overlap with compounds found to be utilized from *Pa* spent media (Supp Table 4). None of the taxa with potential to cross-feed *Pk* were found in the enrichments, probably due to their lower growth rate in sucrose, when compared to *Pa* (Supp Fig 1a, 1b). Overall, these results suggest that when competition for sucrose dominates dynamics, *Pk* will be limited in cross-feeding partners but otherwise may be able to persist by feeding on exudates of diverse bacteria from leaves.

### Richer apoplasts that support more commensal colonization may select against cross-feeders

*Pseudomonas koreensis* (*Pk)* grew on spent medium from *Pantoea* (Pa Fr_-CA_) which had been enriched from *F. robusta* in the S-CA environment. However, *Pa* was also enriched in the S+CA environment where *Pk* could subsist on amino acids (Pa Fr_+CA_) and from *F. trinervia* where it was enriched alone (*Pa* Ft). Growth of *Pk* isolates on spent media of these Pa strains was different. It grew less on exudates of *Pa* enriched from *F. robusta* in the S+CA environment (*Pa* Fr_+CA_20_) and little or none on exudates of *Pa* from *F. trinervia* (*Pa* Ft) (Fig 2c). Corresponding to this growth data, *Pk* Fr_+CA_5_ took up far more metabolites from Pa *Fr* spent media than from Pa *Ft* spent media (Fig 2e, Supp Table 4). We found that inhibitory metabolites cannot explain these differences (Supp Fig 2), so we looked to secreted compounds. The metabolomes of the three Pa strains spent media differed minimally (see CAP analysis in Fig 2d, *p*-value=0.7713) with several compounds previously identified as taken up (hypoxanthine and spermidine) present in all (Supp Table 5). We did identify a few compounds that were taken up uniquely from either of the *Pa* Fr spent media that were not abundant in the other *Pa* strains (Supp Fig 3). These are good metabolite candidates for what drove *Pk* growth, although we could not yet annotate them. Surprisingly, the differences between the *Pa* strains do not correlate to their genetic relatedness: The two Pa Fr strains that support Pk growth shared only ∼81% ANI, but *Pa* Fr_-CA_6_ and the *Pa Ft* isolates share ∼98.6% ANI (see also Supplementary Information and Supp Table 6). Thus, we hypothesized that traits underlying cross-feeding may be selected for in the *Fr* leaf apoplast environment.

The nutrient environment can strongly influence metabolic interactions, so we compared metabolomes of the apoplastic fluid of *F. robusta* (C3 photosynthesis), *F. trinervia* (C4 photosynthesis), and *F. linearis* (uses an “intermediate” C3/C4 photosynthesis). The apoplast metabolomes overall showed a clear separation when considering all detected compounds (Fig 3a). In addition, by targeting amino acids we observed differences in the concentration of methionine, (iso)leucine, valine and lysine, which were higher in *F. trinervia*, while only asparagine was higher in *F. robusta* (Fig 3b and Supp Fig 4). The higher abundance of valuable amino acids in *Ft* also correlated with better apoplast colonization. Two different *Pk* strains successfully colonized *F. robusta* leaves only occasionally, but consistently colonized *F. trinervia* leaves at higher levels (2-sided t-test, p=0.06 and 0.05, respectively, Fig 3c). We did not observe differences in Pk colonization when it was inoculated in *F. robusta* together with *Pa*, but this is likely due to very variable colonization success (Fig 3c). Together, commensal Pk ability to alone colonize *F. trinervia* apoplast, possibly due to availability of resources like important amino acids, could lessen the need for cross-feeding.

**Figure 3.**
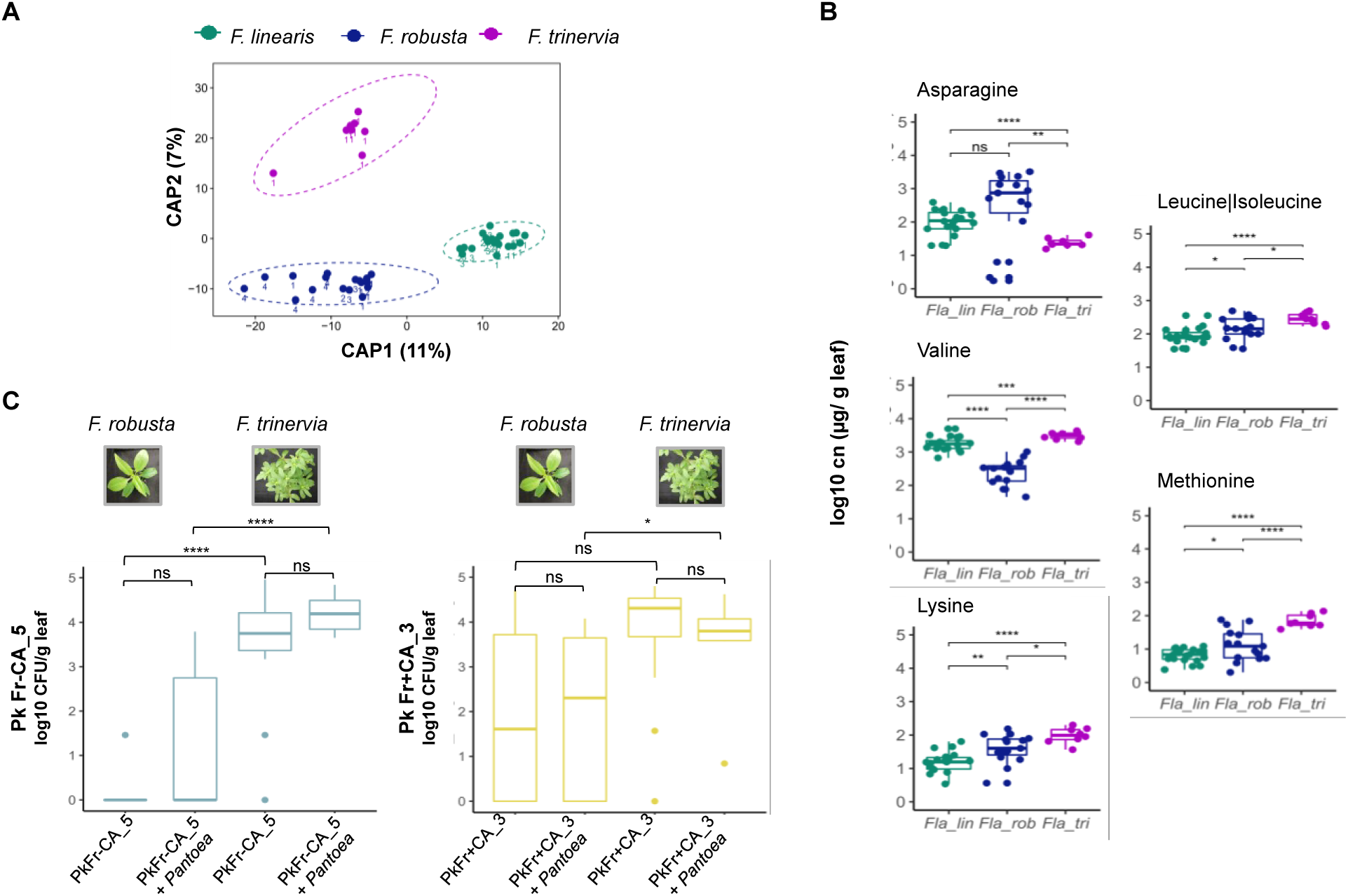
*Pk*Fr-CA_5 requirement of *Pa*Fr-CA_6 when colonizing *F. robusta* might be related to nutrient availability. a) Principle coordinate analysis constrained by plant species, based on all UHPLC-HRMS peaks (area >1.0E4) in untargeted metabolomics of the leaf apoplasts. Dots represent replicate sampling at a given time point, ellipses show 95% confidence intervals, numbers indicate the sampling event; N=20 *F. linearis*, N=18 *F. robusta*, N=8 *F. trinervia*. b) Amino acids showing different concentrations (p-value<0.05) in the apoplast of *F. trinervia and F. robusta*. Concentrations were calculated based on an internal standard. c) Recovery of *Pk* isolates from *F. robusta* and *F. trinervia* leaves 10 days after inoculation, either when inoculated alone or together with the corresponding *Pa*. Left panel shows *Pk* -CA_5 counts and right panel shows *Pk* +CA_3 counts. The experiments were performed twice fully independently, for a total of 8 inoculated leaves of *F. robusta* and 12 of *F. trinervia*.

### Niche differentiation among distinct *P. koreensis* strains maintains diversity during cross-feeding

In the S-CA enrichment, *Pseudomonas koreensis (Pk)* survived exclusively by cross-feeding on diverse resources from *Pantoea* (*Pa)* exudates, but in the S+CA enrichment, *Pk* would have been able to either cross-feed or utilize amino acids or both. Therefore, we hypothesized that multiple *Pk* strains may exist to optimally utilize this niche diversity. Indeed, we observed that the two *Pk* isolates we used *in-planta*, displayed two distinct growth phenotypes on *Pa* spent medium. *Pk* Fr_-CA_5_ grew earlier and reached maximum OD faster in a diauxic pattern compared to *Pk* Fr_+CA_3_ (Fig 4a and Supp Fig 5a). A correlated phenotype was observed in R2 medium, where only the slower cross-feeder switched to more rapid growth when supplemented with vitamins that were present in the enrichment medium (Supp Fig 5b). We predicted that in the S-CA enrichment where cross-feeding was required, faster cross-feeders would outcompete slower ones to dominate the mix, while the more diverse nutrient conditions in the S+CA enrichment would result in a more balanced mix. We checked the phenotype across all *Pk* isolates and surprisingly found that the strains were in similar ratios in both enrichments (Supp Fig 5c, X^2^ p=1). To get an idea of the genetic diversity, we generated draft genomes of the two strains used *in-planta* and a fast grower from the *Fr* S+CA enrichment, *Pk* Fr_+CA_18_. Interestingly, they were all highly similar: all were identified as *Pk* and shared 99.98 to 99.99% ANI (Supp Table 6b). Aligned to the closest reference *Pk* genome, they shared about 97,000 SNPs with only a few hundred SNPs unique to any one genome (Supp Fig 6).

**Figure 4.**
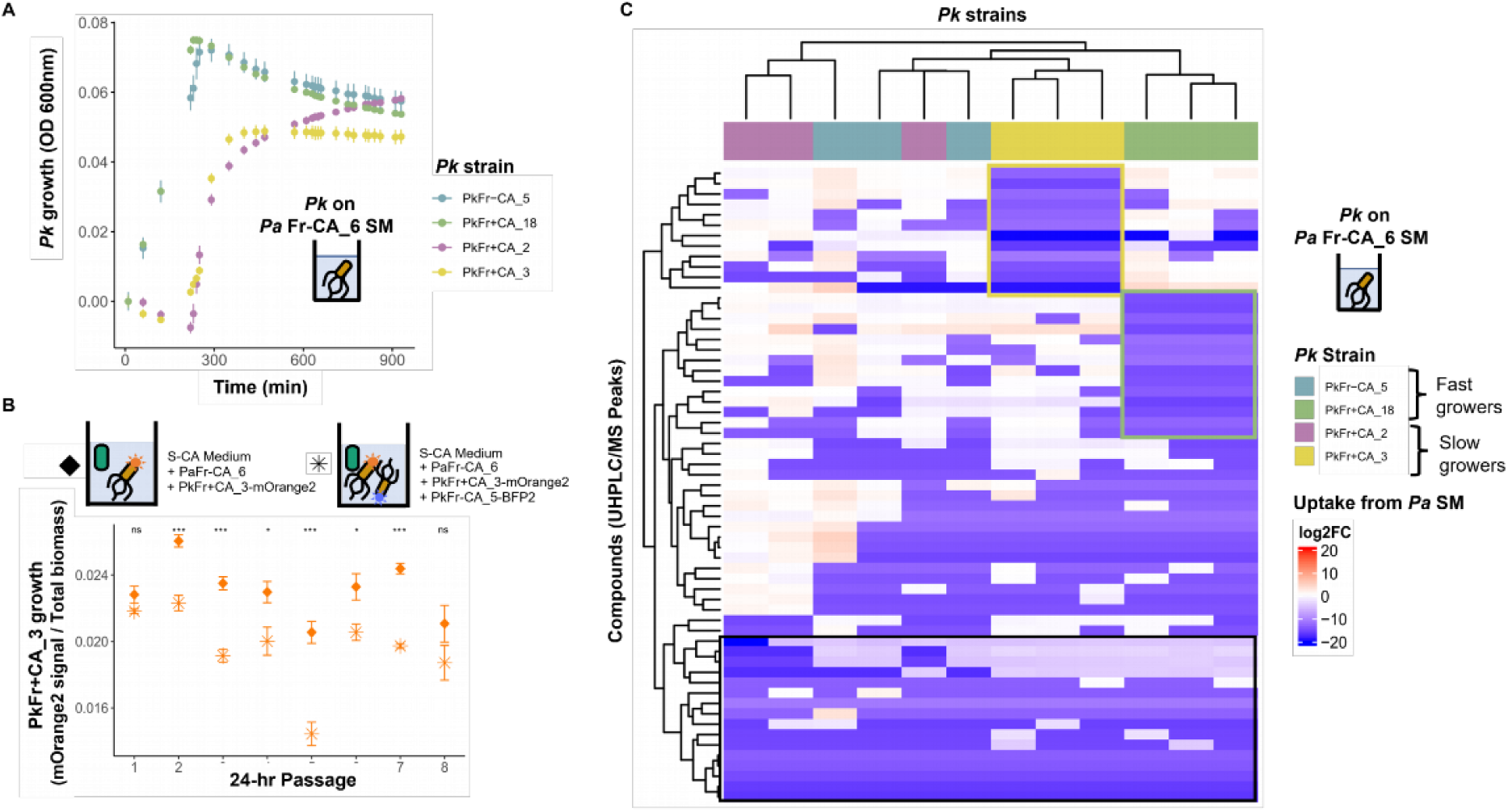
Different *Pk* isolates had specialized niches when feeding from *Pa*Fr-CA_6. A) Growth curves of *Pk*Fr_+CA_ isolates on *Pa*Fr_-CA_6_ spent medium and *Pk*Fr_-CA_5_ for comparison. B) Signal of Pk Fr+CA_3:-mOrange2 fluorescence normalized by the biomass signal in either, co-culture with Pa Fr-CA_6 or in the full community with Pa and Pk Fr-CA-BFP2. C) Consumption of metabolites from *Pa* Fr-CA_6 spent media by different *Pk isolates*. Metabolites shown were significantly taken up (log2 FC <-2, FDR<0.05).

To prove the strains could simultaneously cross-feed from *Pa*, we labeled the slow and fast cross-feeder (*Pk* Fr_+CA_3_ and *Pk* Fr_-CA_5_) with fluorescent tags, which did not alter their growth patterns on *Pa* spent media (Supp Fig 7a). We then combined the strains and grew them directly with *Pa* Fr_-CA_6_ in sucrose-only media. Although the fast cross-feeder was inoculated with only half the cells as the slow cross-feeder (see Supplementary Methods), they were approximately equivalent within 48 hours (Supp Fig 7b). However, neither strain overtook the other after four 48h or eight 24h passages (Supp Fig 7b, 7c). In the 24h passages, we compared growth of the slow grower with *Pa* alone or together with the fast grower. We found that it reached higher levels alone (Fig 4b), consistent with a hypothesis of partially overlapping niches. The fast-growing strain must dominate only some resources, reducing the growth of the slow-grower when they are together, but the slow-grower can persist because it also uses unique resources.

To test this hypothesis, we evaluated growth and uptake of metabolites from *Pa* spent medium by *Pk* Fr_-CA_5_ and *Pk* Fr_+CA_3_ and two additional fast and slow growing isolates (*Pk* Fr_+CA_18_ and *Pk* Fr_+CA_2_, respectively) (Fig 4c). Several metabolite peaks with large negative log2FC (strongest uptake) were shared by all four isolates (Fig 4c and Supp Table 7). Competition for these “preferred” metabolites is consistent with partially overlapping niches. In addition, the isolates significantly depleted unique sets of metabolite peaks (Fig 4c and Supp Table 7). Therefore, our data supports sufficient niche differentiation between the strains to allow very closely related strains, with different rates of growth, to simultaneously cross-feed from *Pa*.

## Discussion

Leaf bacteria play important roles in protecting plants from pathogens and stress via a variety of different mechanisms, so there is broad interest in understanding the factors that shape their abundance (36). It is increasingly clear that interactions between microbial colonizers play major roles in community structuring, whereby the presence of specific microorganisms can drastically alter leaf communities (5). However, the complications of studying microbe-microbe interactions *in-vivo* in leaves severely limits our understanding of the types of interactions that are important. To overcome this, we used *in-vitro* enrichment of leaf bacteria under defined nutrient regimes and could demonstrate that leaf bacteria interact with one another via metabolic cross-feeding. This supports previous work that showed that pervasive cross-feeding is possible between leaf-derived bacteria (37) and that it can explain how a surprising diversity can subsist on single carbon sources. Our work additionally showed that cross-feeding can support bacterial taxa who have no direct utilizable carbon source and that in this case, they can survive only on secreted or leaked metabolites from diverse bacteria.

In previous studies, the diversity of cross-feeding taxa enriched on single carbon sources was high due to promiscuous potential for cross-feeding interactions (37,38). Similarly, we observed that multiple isolated leaf bacterial taxa could have fed *P. koreensis*. However, the cross-feeders in our enrichments consistently were only *Pantoea* and *Pk*. The most likely explanation is simple resource competition. *Pantoea* grew much faster than other isolates on sucrose, allowing it to become the sole feeder of *Pk. Pk* also grew rapidly on *Pantoea* spent medium and could have thereby outcompeted others for key resources. This does not necessarily mean *P. koreensis* would be restricted in cross-feeding partners in leaves. In contrast to the well-mixed and homogeneous *in-vitro* enrichment environment where competition would dominate dynamics, the leaf apoplast is highly compartmentalized and heterogeneous (39). Additionally, most endophytic commensal bacteria in leaves reach only low colonization density (40). These factors would decrease the importance of resource competition, so that diverse taxa can establish a niche consuming primary plant-derived resources and *Pk* or others could take up leaked metabolites.

Technological barriers currently limit our understanding of how often cross-feeding really occurs in leaves and its benefit to plants. However, recent improvements to inter-bacterial interaction inference suggests positive interactions have probably been underestimated (40) and experimental results suggest frequent positive interactions among co-colonizing leaf bacteria (41,42). This might seem unlikely, since ecological models have predicted that cooperative interactions can destabilize microbial communities and that competition should therefore benefit hosts (43). However, the instability in these models is caused by strong species dependencies that can easily be interrupted. In contrast, cross-feeding among leaf bacteria seems to involve weak coupling and high promiscuity. This type of cross-feeding can actually stabilize microbiomes to invasion, because redundant cross-feeding networks leave few resources for invaders (44). Apoplast nutrients are important regulators of *Pseudomonas syringae* virulence, so full occupation of resources by cross-feeding could also limit damage caused by pathogens (45). This could explain how *Pantoea* protects crops from pathogenic *P. syringae* by decreasing its virulence without eliminating it from the leaf microbiome (46). Additionally, cross-feeding would be beneficial if it allows bacteria to establish that can switch to competitive behaviors upon invasion that protect plants, such as antibiotic production (47). Therefore, more thorough investigations into the role of cross-feeding in interactions between leaf bacteria and pathogens and its general role in shaping and stabilizing co-colonizing leaf microbiota are needed.

The four *Pseudomonas koreensis* isolates we sequenced were genetically similar with ANI of ∼99.98% to 99.99% and ∼97,000 shared SNPs against the most similar *P. koreensis* reference genome. While more study is needed to understand the arisal of this diversity, it appears to be functionally relevant. The sequenced strains had distinct cross-feeding niches with only partially overlapping metabolite uptake profiles, making it possible to cross-feed in parallel despite different growth rates. This diversity must have originated in the original leaf communities since the phenotypes appeared both in sucrose-only enrichments where cross-feeding was required and in enrichments with amino acids as additional resources. Similar levels of diversity were previously found in leaf-associated *Pseudomonas viridiflava*, where a single OTU (99% 16S rRNA gene sequence similarity) harbored at least 82 distinct strains (99.9% genome identity), with clearly different host interaction phenotypes (8). Commensal *Sphingomonas* bacteria were also shown to exhibit extensive diversity in the presence of secretion systems likely relevant for microbe-microbe interactions (48). Our results extend the functional relevance of leaf bacterial diversity to inter-bacterial metabolic interactions. Given that nutrition plays key roles in virulence (49,50), it also raises the intriguing question of whether interactions like cross-feeding may ultimately influence interactions with hosts.

*Pantoea* isolates also exhibited interesting and surprising functional diversity. Those from *F. robusta* enrichments, where cross-feeding occurred, fed *Pseudomonas* better than those from *F. trinervia* enrichments, probably by producing more of some key metabolites. Surprisingly, the “good” cross-feeders enriched from *F. robusta* were only distantly related (81% ANI), consistent with inter-species differences (51) compared to 98-99% ANI between good and bad cross-feeders. This result is consistent with the *F. robusta* environment selecting for *Pantoea* with increased metabolite secretion, benefitting cross-feeders. Both *Pantoea* and *P. koreensis* persisted in *F. trinervia* leaves alone at higher levels than *F. robusta*. This could be due to relatively lower levels of key nutrients. Indeed, we found lower levels of methionine, isoleucine, lysine and valine in *F. robusta*. Except for the aromatic amino acids, methionine, isoleucine and lysine are three of the four biosynthetically highest-cost amino acids (together with histidine, (52). Additionally, we observed cross-feeding purines guanine and hypoxanthine, which are connected to methionine via the THF cycle. Thus, differences in the apoplast metabolite landscape could alter selection for traits underlying cross-feeding. In extreme cases, increased metabolite secretion could result from evolution of reciprocal cross-feeding between *Pantoea* and *Pseudomonas* specifically, which can in turn lead to strong co-adaptation (23). However, this seems unlikely in leaves since conditions including compartmentalization, low bacterial density and high bacterial diversity are likely to make interactions between specific taxa transient, decreasing the likelihood that reciprocal cross-feeding will arise (53). Thus, if more metabolite secretion is a beneficial adaptation in the *F. robusta* apoplast environment, it is more likely because *Pantoea* derives benefits from cross-feeding with diverse taxa rather than *Pk* alone.

The apoplast is together with the roots a critical site of host interaction with microbes. While it is not yet clear why some amino acids and other metabolites differ strongly between *Flaveria* species, one possibility is their different photosynthesis mechanisms. Evolution of C4 photosynthesis in *Flaveria* has had diverse effects. For example, higher glutathione turnover in sulfate assimilation (28) has led to higher cysteine levels in *F. trinervia* leaves. While we could not reliably quantify cysteine, it is the direct precursor to methionine, which was significantly elevated in *F. trinervia* apoplast. Thus, links between photosynthesis and apoplast metabolites are plausible. This is reminiscent of roots, where the exudate nutrient landscape key roles in shaping microbial communities (54,55) and differs between different plant species (56,57). Our results strongly suggest that apoplast exudates influence the arisal of microbial interactions, which in turn help shape leaf microbiomes (5). If so, this is an exciting prospect because it could offer targets for manipulation by plant breeders.

## Acknowledgements

We wish to thank Carl-Eric Wegner, Stefan Riedel and Professor Kirsten Küsel for making their sequencing equipment and knowledge available to us. We gratefully acknowledge the Deutsche Forschungsgemeinschaft CRC 1127 “ChemBioSys” who funded the UHPLC-HRMS system used for metabolomics experiments here. MTA is supported by the Carl Zeiss Stiftung via the Jena School for Microbial Communication and were funded by the Deutsche Forschungsgemeinschaft (DFG, German Research Foundation) under Germany’s Excellence Strategy – EXC 2051 – Project-ID 390713860. MMR is supported by the International Leibniz Research School.

## Data availability

Supplementary methods, figures and tables are provided in the files SI and SI Tables 2,4,5, and 7. Scripts and data used to recreate all figures are publicly available via Figshare: https://figshare.com/projects/Niche_separation_in_cross-feeding_sustains_bacterial_strain_diversity_across_nutrient_environments_and_may_increase_chances_for_survival_in_nutrient-limited_leaf_apoplasts/125920.

Raw sequencing data has been made publicly available in the NCBI project PRJNA778092 and raw metabolomic data is being made publicly available in MetaboLights under the study MTBLS3719.

