## Supplemental Information for "Niche separation in cross-feeding sustains bacterial strain diversity across nutrient environments and may increase chances for survival in nutrient-limited leaf apoplasts"

Contents:

Supp Figures 1-7
Supp Tables 1,6,8 and 9
Supplementary Methods

Supplementary Figure 1

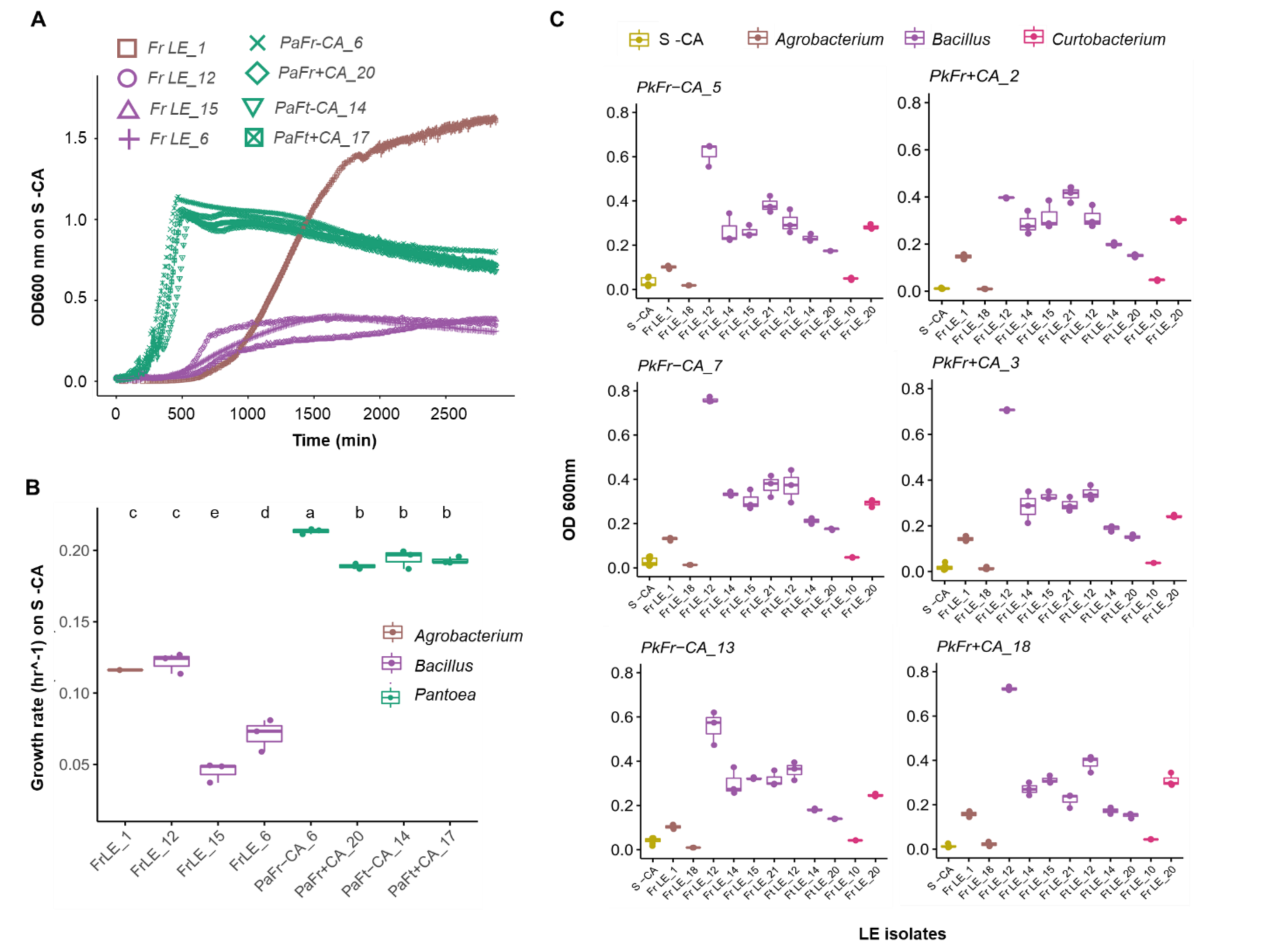


**Supplementary Figure 1. Diverse isolates could cross-feed *Pk* but they were probably outcompeted on sucrose by *Pa.*** A) Growth in S -CA medium of different *Agrobacterium* and *Bacillus* isolates versus the *Pa* isolates. B) Growth rates in S -CA medium of the same isolates as in panel A. C) Growth of several *Pk* isolates on spent media from different *Agrobacterium, Bacillus* and *Curtobacterium* isolates obtained from the leaf extracts of *F. robusta* and *F. trinervia*.

Supplementary Figure 2

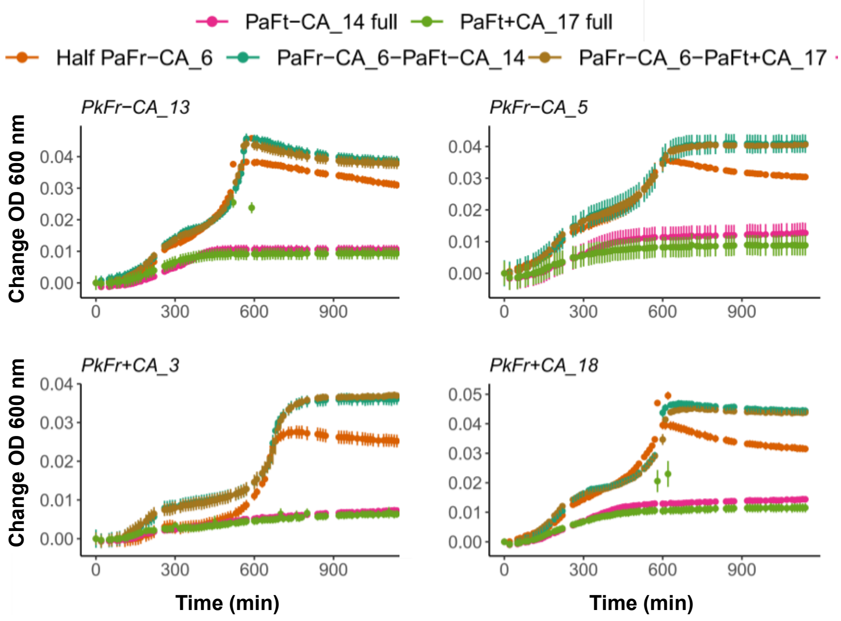


**Supplementary Figure 2**. **Low growth of *Pk* isolates on *Pa*Ft spent media is not due to inhibition.** Growth of several *Pk* Fr isolates on *Pa* Ft spent media, either at full concentration, or in 1:2 combination with spent media of *Pa* Fr-CA_6.

Supplementary Figure 3

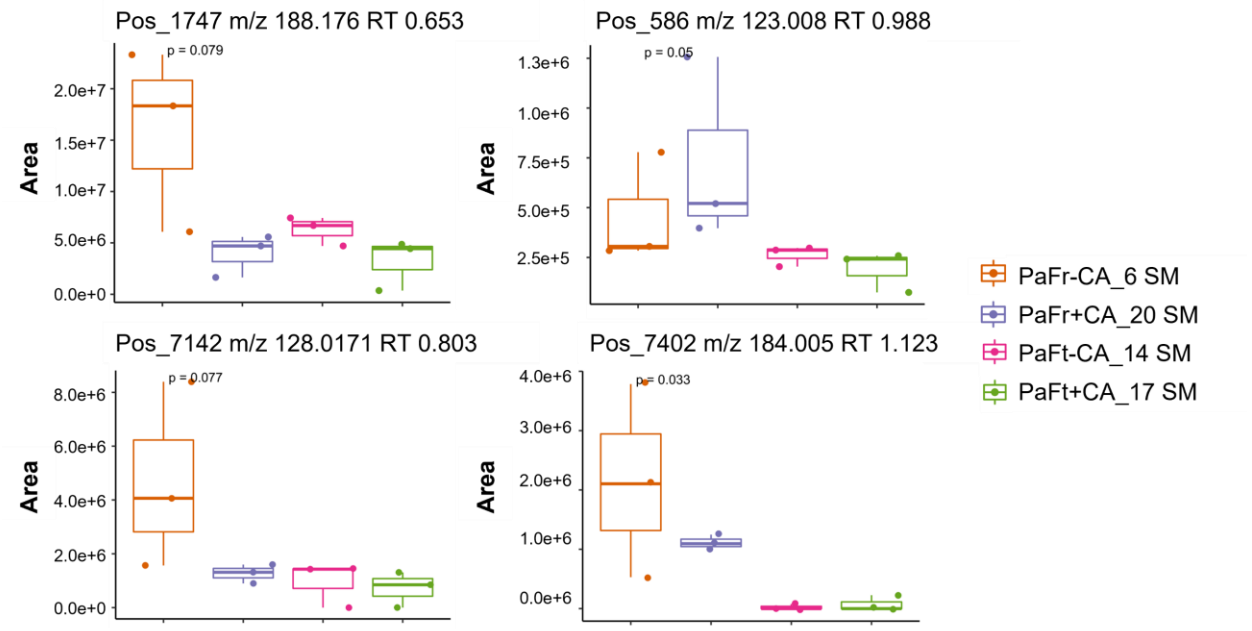


**Supplementary Figure 3. Areas detected in the different Pa spent media of compounds commonly taken up from the two *good Pantoea* Fr cross-feeders.** For each peak, the ionization mode in which it was detected (Pos=positive, Neg=negative), the m/z value and the retention time are shown. P-values are based on Kruskal-Wallis test.

Supplementary Figure 4

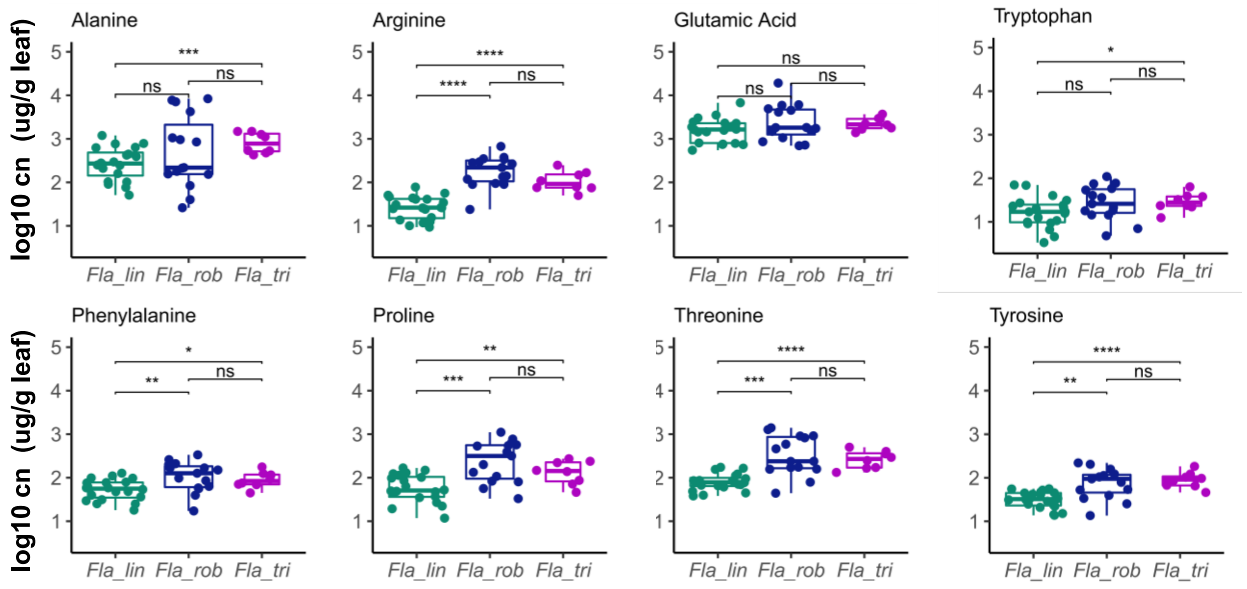


**Supplementary Figure 4. Amino acid concentration in the apoplast of *Flaveria* leaves.** N=20 *F. linearis,* N=18 *F. robusta, N*=8 *F. trinervia.* Concentrations were calculated based on the deuterated amino acid internal standard.

Supplementary Figure 5

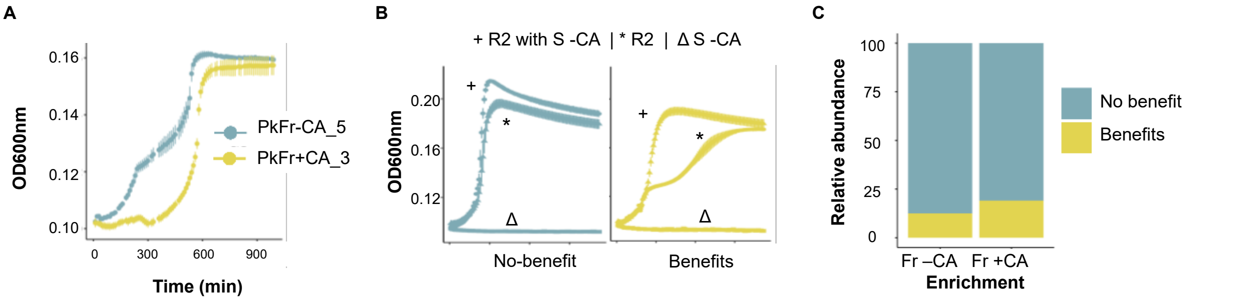


**Supplementary Figure 5. The enrichments harbored metabolically diverse *Pk* isolates with different nutrient dependencies.** A) Growth curve of a fast (*Pk* $Fr_{-CA\_5}$) and a slow (*Pk* $Fr_{+CA\_3}$) grower isolate on *Pa*$Fr_{-CA\_6}$ spent medium. B) Effect of nutrient supplementation on the growth of the two isolates. C) Distribution of the two growth-phenotypes across the *Pk* isolates of each enrichment.

Supplementary Figure 6

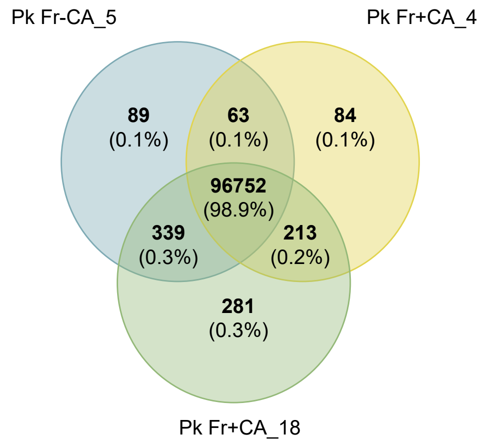


**Supplementary Figure 6. *Pk* isolates with distinct metabolic phenotypes have extremely similar genomes.** A) Venn Diagram of single nucleotide variants across three *Pk* isolates after annotation with the reference genome of *P. koreensis* D26.

Supplementary Figure 7

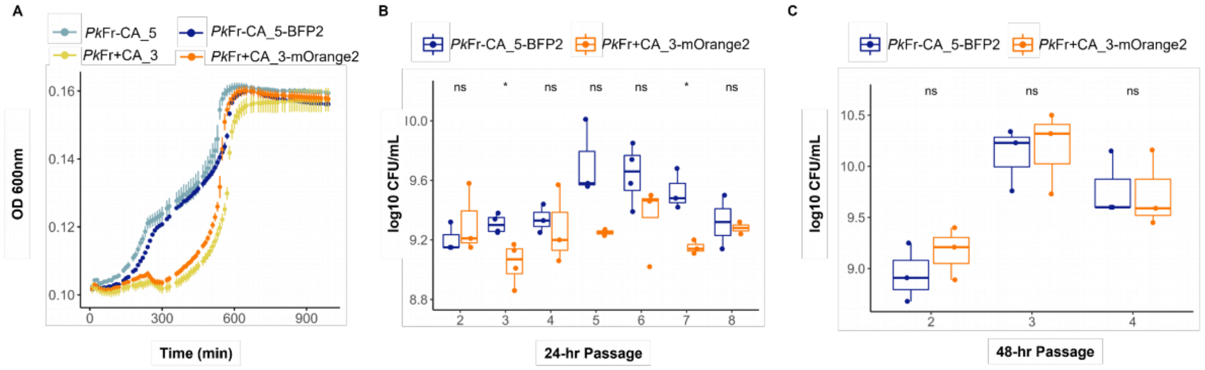


**Supplementary Figure 7. Tagging did not affect the strains phenotypes.** A) Growth of tagged versus non-tagged *Pk* isolates on *Pa* Fr-CA_6 spent media. ***Pk*Fr-CA_5 dominated the full community but did not outcompete *Pk*Fr+CA_3.** B) CFU counts of each tagged *Pk* in the full community passaged every 24 hours. C) CFU counts of each tagged *Pk* in the full community passaged every 48 hours.

**Supplementary Table 1**. Volume (µL) of bacterial suspension (tagged *Pk* isolates and *Pa*Fr$_{-CA_6}$) and PBS added in each condition for the co-culture assay.

| **Condition** | ***Pa*Fr$_{-CA_6}$** | ***Pk*Fr$_{-CA_5:BFP2}$** | ***Pk*Fr$_{+CA_3:mOrange2}$** | **1x PBS** |
|---|---|---|---|---|
| *Pa*Fr$_{-CA_6}$ + *Pk*Fr$_{-CA_5:BFP2}$ | 10 | 3.3 | --- | 6.6 |
| *Pa*Fr$_{-CA_6}$ + *Pk*Fr$_{+CA_3:mOrange2}$ | 10 | --- | 6.6 | 3.3 |
| *Pa*Fr$_{-CA_6}$ + *Pk*Fr$_{-CA_5:BFP2}$ + *Pk*Fr$_{+CA_3:mOrange2}$ | 10 | 3.3 | 6.6 | --- |
| *Pa*Fr$_{-CA_6}$ | 10 | --- | --- | 9.9 |

**Supplementary Table 6.**

A) Average nucleotide identity (%) between the genomes of several *Pa* isolates.

| | **Pa Ft$_{-CA_14}$** | **Pa Fr$_{+CA_20}$** | **Pa Ft$_{+CA_17}$** | **Pa Fr$_{-CA_6}$** |
|---|---|---|---|---|
| **Pa Ft$_{-CA_14}$** | | 81.160 | 99.997 | 98.685 |
| **Pa Fr$_{+CA_20}$** | 81.160 | | 81.292 | 81.207 |
| **Pa Ft$_{+CA_17}$** | 99.997 | 81.292 | | 98.678 |
| **Pa Fr$_{-CA_6}$** | 98.685 | 81.207 | 98.678 | |

B) Average nucleotide identity (%) between the genomes of several *Pk* isolates

| | **Pk Fr$_{-CA_5}$** | **Pk Fr$_{+CA_3}$** | **Pk Fr$_{+CA_18}$** |
|---|---|---|---|
| **Pk Fr$_{-CA_5}$** | | 99.9909 | 99.9906 |
| **Pk Fr$_{+CA_3}$** | 99.9909 | | 99.9841 |
| **Pk Fr$_{+CA_18}$** | 99.9906 | 99.9841 | |

**Supplementary Table 8**. Oligonucleotides used in this study.

| Name | Target | Sequence 5' to 3' * | References |
|---|---|---|---|
| B341F | 16S rRNA V3/V4 | CCTACGGGNGGCWGCAG | (Muyzer et al., 1993)/(Caporaso et al., 2010) |
| B806R | 16S rRNA V3/V4 | GGACTACHVGGGTWTCTAAT | |
| FWD Tn*5/7*_gt | Veryfing the Tn7 insertion | ATGGTGAGCAAGGGCGAG | (Schlechter et al., 2018) |
| REV_Tn*5/7*_gt | Veryfing the Tn7 insertion | CAACAGGAGTCCAAGCTCAG | |
| Tn*7*_gt1 | Genotyping the Tn7 backbone | GAATTACAACAGTACTGCGATGAG | |
| Tn*7*_gt2 | Genotyping the Tn7 backbone | GATCAACTCTATTTCTCGCGGG | |
| Tn*7*_gt3 | Genotyping the Tn7 backbone | TACATAACGGACTAAGAAAAACACTACAC | |
| FWD_uidA | *uidA* gene (*E. coli*) | AACAGGTGGTTGCAACTGGA | |
| REV_uidA | *uidA* gene (*E. coli*) | TTGCTGAGTTTCCCCGTTGA | |

**Supplementary Table 9.** Concentration of deuterated amino acids in CELL FREE AMINO ACID MIX (U-D, 98%), from Cambridge Isotope Laboratories, Inc. (MA, USA). Lot# PR-29964

| Deuterated amino acid | Weight percentage (m/m) |
|---|---|
| Alanine_D | 6.80% |
| Arginine_D | 3.10% |
| Asparagine_D | 4.40% |
| Aspartate_D | 4.90% |
| Glutamine_D | 4.50% |
| Glutamic Acid_D | 10.30% |
| Histidine_D | 1.20% |
| Lysine_D | 8.70% |
| Leucine_D\|Isoleucine_D | 18.90% |
| Methionine_D | 1.90% |
| Phenylalanine_D | 6.30% |
| Proline_D | 2.60% |
| Serine_D | 3.80% |
| Threonine_D | 5.10% |
| Tryptophan_D | 2.60% |
| Tyrosine_D | 3.00% |
| Valine_D | 6.50% |

**Supplementary Methods**

Enrichment of leaf microbiomes from *Flaveria trinervia* and *Flaveria robusta*

Cuttings from *Flaveria robusta, F. linearis* and *F. trinervia* were grown in pots containing a mixture of soil (66%) perlite (33%), and Substral Osmocote® NPK (Mg) 17-9-11 (2) (4 g/L of soil mixture). The plants were kept at an average day/night temperature of 25 °C/22 °C and a photoperiod of 16 hours in our lab. After roots were developed (~3 weeks), they were transplanted into an outdoor garden (Jena, Germany) to allow natural colonization by microorganisms. After two months, well-developed leaves from both species were sampled, weighed and washed three times in sterile water to remove dirt and insects. Leaf extracts were prepared by macerating the washed leaves with a sterile pistil, adding 1 mL of 1X PBS with 0.02% Silwet and vortexing for 10 seconds, followed by quick centrifugation to precipitate down the larger segments. The supernatant was transferred to a clean tube, mixed with glycerol to a final 20% v/v concentration and stored at -80 °C. The number of live bacterial cells were estimated by plating serial dilutions on Reasoner's 2A agar (R2A) and calculating CFU/mL.

Next, the leaf extracts were enriched *in-vitro* over 12 passages on two different sucrose-based minimal media. First, a preculture was generating by inoculating ~1000 cells in 15 mL of basic M9 broth (2 mM MgSO4*7H20, 0.2 mM CaCl2, 1 µg/mL biotin, 1 µg/mL thiamine, 11 mM sucrose and trace elements: 134 µM EDTA, 31 µM $FeCl_3$-$6H_2O$, 6.2 µM $ZnCl_2$, 0.76 µM $CuCl_2$-2H2O, 0.42 µM $CoCl_2$-2H2O, 1.62 µM $H_3BO_3$, 0.081 µM $MnCl_2$-$4H_2O$), supplemented with 0.2% w/v casamino acids (Difco) 200 mM $NH_4Cl$ and 200 µg/mL of cycloheximide to limit eukaryotic growth. The cultures were incubated at 26 °C and 220 rpm for 72 hours. One fraction of the culture was used to prepare 20% glycerol stocks. The remaining volume was centrifuged (5000 x g for 5 min), the pellet washed twice in 1X PBS, and resuspended to a final $OD_{600nm}$ of 0.3 in fresh 1x PBS. Five microliters of the suspension were inoculated in a 2 mL 96 well plate containing 1 mL M9 media supplemented with $NH_4Cl$ (33 mM) and either no casamino acids (S -CA) or 0.2% m/v casamino acids (S +CA). Each enrichment condition was run in triplicates. The plate was incubated at 26 °C and 220 rpm. Every 48 hours, the cultures were homogenized by pipetting up and down and 5 µL of each well were transferred to a new plate with fresh media. The $OD_{600nm}$ was measured at each passage, and the procedure was repeated 12 times. In the first two passages, cycloheximide was added in the same concentration as before. At the last passage, 700 µL of each well were pelleted for DNA extraction by centrifuging at 20000 x g for 10 min. The remaining volume of the three replicates were combined to prepare 20% glycerol stocks, which were stored at −80 °C.

Characterization of culture-independent bacterial diversity in enrichments

The bacterial communities in the twelfth passage were characterized by 16S rRNA gene amplicon sequencing. Cell pellets were resuspended in 600 µL SDS extraction buffer (SDS extraction buffer, 10% filter sterilized SDS, 100 mM Tris pH 8.0, 200 mM NaCl, 2 mM EDTA), and transferred to 2 mL screw-cap tubes containing ~0.2 g of glass beads (0.25-0.50 mm, ROTH). The mixture was incubated at 37°C for 10 min, followed by bead beating for 30 sec at 1400 rpm in a BioSpec Mini-Beadbeater-96 and 4 minutes centrifugation at 13400 rpm. The supernatant was transferred into new 1.5 mL sterile

Eppendorf tubes, and 200 µL of 5 M potassium acetate were added to precipitate the SDS. After 5 minutes of centrifugation at 13400 rpm, the supernatant was recovered in a clean tube. The DNA was purified using 1.5x volume of Sera-Mag™ magnetic carboxylate modified particles and eluted in 50 µL of 10 mM TrisHCl (pH 8.0). The purified DNA was stored at −20 °C further amplification. Amplification of the 16S V3-V4 region was carried out in a two-step PCR; on the first step, the samples were amplified using the bacterial primers B341F and B806R(Caporaso et al., 2010; Muyzer et al., 1993) (Supp Table 8). The PCR mastermix contained: 8 µL KAPA 5x GC buffer, 0.3 mM KAPA DNTPs, 0.8 µL Kapa HiFi polymerase (KAPA Biosystems), 0.075 µM of each primer, 1 µL of template and 28.4 µL NFW. The amplification settings were 2 min denaturing at 95 °C, followed by 15 cycles of denaturation at 98 °C for 30 seconds, annealing at 50 °C for 30 s, and elongation at 72 ° C for 40 sec, followed by a final extension at 72 °C for 2 minutes. The PCR products were enzymatically cleaned with Antarctic phosphatase and Exonuclease I (New England Biolabs, Inc) (0.5 µL each enzyme with 1.22 µL Antarctic phosphatase buffer at 37 °C for 30 minutes followed by 80 °C for 15 min). In the second step, concatenated primers were used; these were designed to include an Illumina adapter P5 (forward) or P7 (reverse), an index sequence, a linker region, and the B341F and B806R primers. Each sample was amplified in triplicates; the Mastermix consisted of 3 µL KAPA 5x GC buffer, 0.3 µM KAPA DNTPs, 0.3 µL KAPA HiFi polymerase (KAPA Biosystems), 0.015 µM of each primer, 1 µL of the cleaned product from the first PCR and 8.25 µL of NFW. The cycling settings were: 2 min denaturing at 95 °C, followed by 25 cycles of denaturation at 98 °C for 30 seconds, annealing at 50 °C for 30 s, and elongation at 72 ° C for 40 sec, followed by a final extension at 72 °C for 2 minutes. Samples that did not amplify with these settings were amplified with 30 cycles instead. Triplicate reactions were pooled and purified with 0.65x volume magnetic beads and eluted in 10 mM TrisHCl and run in a 1.5% agarose gel. The brightness of each band in the gel was measured in ImageJ (version 1.52a) and used to adjust the volume of each sample in the final library. After combining all samples, the pool was purified once again with 0.8x volume Sera-Mag beads and quantified in a Qubit (Thermo Fisher Scientific, Inc). The library was denatured and then loaded onto a MiSeq lane spiked with 10% PhiX genomic DNA to ensure high sequence diversity and sequenced in 500 cycles (2x250 bp).
Amplicon sequencing data was split on the indices and the adapters were trimmed from the read ends using Cutadapt. The data was clustered into amplicon sequencing variants “ASVs” using dada2, setting the quality filter at (truncLen=c(200,200)). Next, the sequences were dereplicated to remove redundant reads, followed by denoising using the error rate and calling of ASVs in the forward and reverse reads. When merging the forward and reverse reads, only overlapping regions were kept. Chimeric sequences were removed and obtained a sequence table with the merged data. For taxonomy assignment, the Silva database (v132) was used. Data analyses were carried out in R (version 4.0.4) with the packages *phyloseq* and *vegan.*

Characterization of culturable bacterial diversity in original leaf extracts and in enrichments

Isolates were recovered from the initial leaf extract glycerol stocks and from the glycerol stocks from the twelfth enrichment passage of each condition. Serial dilutions from the glycerol stocks were prepared in 1x PBS and spread in R2A plates. From each condition, 25 random isolates were selected and restreaked to recover pure cultures (150 isolates total). To minimize isolate adaptation to plating, no further cultivation was performed and pure isolates were grown in R2 broth for 48 hours to prepare 20% glycerol stocks. All isolates were identified via DNA extraction and Sanger sequencing of the 16S rRNA gene. Liquid cultures in R2 broth of each isolate were pelleted by centrifugation at 20000 x g for 10 min and stored at -20 °C. DNA extraction and purification was done as described in the previous section. Whole 16S rRNA gene was amplified with the universal primers 8F and 1492R (Supp Table 8). The PCR mastermix contained: 8 µL KAPA 5x GC buffer, 0.32 µM KAPA DNTPs, 0.8 µL KAPA HiFi polymerase (KAPA Biosystems), 0.27 µM of each primer, 1 µL template and 24.2 µL nuclease free water (NFW). The amplification consisted of 2 min denaturing at 95 °C, followed by 30 cycles of denaturation at 95 °C for 30 seconds, annealing at 50 °C for 30 s, and elongation at 72 ° C for 1:30 min, followed by a final extension at 72 °C for 5 minutes. and the products were Sanger sequenced (Eurofins Genomics, Germany). The sequences were trimmed in R (version 4.0.4) using the package *sangeranalyseR* with default parameters. The resulting fasta file was "BLASTed" against the NCBI 16S rRNA gene database. The first ten hits were used to identify taxonomy at the genus level using a least common ancestor algorithm in MEGAN6 with percent to cover set to 70 and the minimum percent identity to 97 (Huson et al., 2016).

Whole Genome sequencing

Whole genome sequencing was carried on three *Pseudomonas koreensis* (*Pk*Fr$_{-CA_5}$, *Pk*Fr$_{+CA_3}$ and *Pk*Fr$_{+CA_18}$) and four *Pantoea sp.* isolates (*Pa*Fr$_{-CA_6}$, *Pa*Fr$_{+CA_20}$, *Pa*Ft$_{-CA_14}$, and *Pa*Fr$_{+CA_17}$). For this, a modified DNA extraction protocol was used. The isolates were grown overnight in 2 mL of R2 broth at 28 °C and 220 rpm. The cultures were pelleted by centrifugation (20000 xg for 5 min), resuspended in 600 µL SDS extraction buffer and transferred to 2 mL screw-cap tubes containing ~0.2 g of glass beads (0.25-0.50 mm, ROTH). The mixture was incubated at 37°C for 10 min, followed by bead beating for 30 sec at 1400 rpm in a BioSpec Mini-Beadbeater-96 and five minutes centrifugation at 20000 x g. The supernatant was transferred into new 1.5 mL sterile Eppendorf tubes to which 100 µg/mL of Proteinase K (Sigma) were added. The mixture was incubated for one hour at 37 °C, followed by 10 min at 80 °C to deactivate the enzyme. Once the tube had cooled, 10 µg/mL RNAse A was added, and incubation was carried at 37 °C for 30 min. To remove proteins, an equal volume of phenol: chloroform: isoamyl alcohol (25:24:1) were added to each sample, followed by 10 min centrifugation at 4 °C and 20000 x g. The top layer was transferred to a clean tube where an equal volume of chloroform / isoamyl alcohol (24:1) was added. The centrifugation step was repeated, and again the top layer was transferred to a new tube. One-tenth volume of 3M sodium acetate and 2.5 volumes of 100% ethanol were added, and the tubes were left overnight at -20 °C. To pellet the DNA, the tubes were spun for 40 minutes at 20000 x g and 4 °C; the supernatant was removed, and the pellet was washed with 200 µL of 70% ethanol and left to air dry. Finally, the DNA was resuspended in 100 µL of 10 mM TrisHCl. The samples were sent to Microbial Genome Sequencing Center (Pittsburgh, USA) for sequencing on the NextSeq 2000 platform at a depth of 300 MBp (~50x for P. koreensis strains and ~60x for P. agglomerans strains).

Whole genome assembly, annotation, ANI calculation and single nucleotide polymorphism detection

For each isolate, the raw forward and reverse reads were trimmed using Trimmomatic (version 0.39), which removed the Illumina adapters, and trimmed at quality below 15 in a 4-base wide sliding window. The assembly of the genomes was done using SPADES (3.14.1) with the default parameters in “isolate” mode. Average nucleotide identity between the different isolates of each genera was calculated in Kbase (Arkin et al., 2018). First, the final scaffolds from the SPADES assembly were imported into Kbase and were annotated using Prokka (v1.12). The average nucleotide identity was then calculated using the FastANI app.
Single nucleotide polymorphisms (SNPs) compared to the *Pseudomonas koreensis* D26 reference genome (assembly accession GCF_001605965.1) were called by mapping the raw sequencing reads from each *P. koreensis* strain using SNIPPY (version 4.6.0). A custom script was then used to parse the resultant SNP files to determine which SNPs were shared or unique for each *P. koreensis* isolate.

Evaluation of the isolates carbon preference

To test metabolic dependencies among the communities, we grew all 150 isolates individually in S -CA and S +CA media. Single colonies taken from R2A plates were inoculated in triplicates in a 96-well flat bottom plate, containing 200 µL of media per well. The plates were incubated at 26 °C and 220 rpm in a shaker for 24 h. The $OD_{600}$ was measured right after inoculation and after 24 h. A second passage over the same media was done to verify the observed growth was not due to carryover of R2A nutrients. An isolate was categorized as growing if its final OD was ≥0.1. Of the isolates that showed growth without CA supplementation, we chose several and measured their growth rate in the S -CA medium. For this, the isolates were pre grown in S -CA broth for 24 hours at 220 rpm and 28 °C. The cells were washed twice with 1X PBS, centrifuging each time at 5000 rpm for 5 min and discarding the supernatant in between. The washed cells were resuspended in 1X PBS and diluted to an $OD_{600nm}$ of 0.2. In a 96-well plate, 20 µL of the bacterial suspension were mixed with 180 µL of S -CA broth. All strains were inoculated in triplicates. The plate was incubated in a VersaMax Tunable Microplate Reader (Molecular Devices, CA) for 48 hours at 28 °C, taking $OD_{600nm}$ measurements every 10 min and mixing in between reads. To determine the growth rate of each curve, we calculated the first derivative in 30-min windows across the length of the curve, then we selected the periods where the derivative was larger than the average variation (i.e., the exponential phase), and calculated the mean derivative for those time points. Several isolates that could not grow without CA were tested for amino acid auxotrophy on a modified version of the S -CA media, where sucrose was replaced by 22 mM glucose. Additionally, we checked whether they could consume sucrose in presence of other nutrients. For this, they were precultured in R2 broth overnight at 28 °C and 220 rpm, harvested and diluted to an OD of 0.2 as detailed above and inoculated in a 96-well plate containing 180 µL of each of the following S -CA: R2 broth: sterile water ratios; 1:0:0.8, 1:0.2:0.6, 0:0.2:1.6. Each strain was inoculated in triplicates in each of the five conditions by adding 20 µL of the cell suspension (final OD 0.02). The plate was incubated in the plate reader with the same settings as before.

Tagging of *Pseudomonas* strains with fluorescent proteins and competition assay

To test whether faster growing *Pseudomonas* isolates would outcompete slow growing isolates when growing along with *Pantoea*, we tagged the isolates PkFr$_{-CA_5}$ and PkFr$_{+CA_3}$ with the fluorophores mTagBFP2 and mOrange2 respectively, using the delivery plasmid systems pMRE-Tn7-140 and pMRE-Tn7-144 developed by Schlechter et al. (2018). In brief, *E. coli* ST18 containing each plasmid was grown overnight in LB broth + 5-aminolevulinic acid (ALA) (50 µg/mL) + Amp (100 µg/mL) + Gen (15 µg/mL) + Cam (15 µg/mL) at 30°C and 200 rpm. The *Pseudomonas* strains were grown in LB without ALA or antibiotics under the same incubation settings. On the following day, 25 mL of LB + ALA + antibiotics were inoculated with 500 µL of the overnight *E. coli* cultures and 25 mL of LB without antibiotics were inoculated with 1000 µL of the *Pseudomonas* culture and grown until an $OD_{600nm}$ of about 0.7. The cells were harvested by mild centrifugation (2000 x g for 5 min), and after discarding the supernatant, they were resuspended in 1X PBS by gently inverting the tubes. The *Pseudomonas* and the corresponding *E. coli* cultures were mixed in three different ratios; 1:3, 1:1 and 3:1 considering the $OD_{600nm}$ of the suspensions. The combined cells were harvested as before, resuspended in 100 µL of 1X PBS and drop spotted in an LB + ALA plate containing 100 µL 1 M CaCl2 to increase conjugation efficiency. The plates were incubated at 30 °C overnight; the next day a colony was suspended in 1 mL 1X PBS and different volumes (100-300 µL) were spread in LB plates containing antibiotics and incubated at 30 °C for several days. Colonies that appeared in the plates were checked for fluorescence under UV light (330 nm) and in a Axiozoom Stereomicroscope (Zeiss). To rule out the selected colonies were not *E. coli*, a PCR using the specific primers FWD_uidA and REV_uidA (Supp Table 8) was performed. Additionally, the presence of the fluorescent protein coding sequence was checked with the primers FWD_Tn5/7_gt and REV_Tn5/7_g (Supp Table 8). In both cases, the Mastermix consisted of: 1x Buffer B (Biodeal), 0.4 mM dNTPs (Carl Roth), 0.2 µM of each primer, 2.5 mM $MgCl_2$ and 0.25 µL Taq DNA polymerase (Biodeal). For the *E. coli* specific primers, the cycling conditions were: 3 min at 95 °C, 30 cycles of 30 sec at 95°C, 30 sec at 57.5 °C and 1 min at 72 °C, followed by a 5 min final extension at 72 °C. For the set of primers targeting the transposon, the cycling was the same, except for the annealing temperature, which was set at 56.5 °C. Once the colonies were confirmed to be fluorescent *Pseudomonas*, they were grown in liquid LB + Gen + Cam at 37 °C and 200 rpm to cure them from the plasmid. The efficacy of this last step was confirmed via PCR with the primers FWD_Tn7_gt, Rev_Tn7_gt and Tn7_gt (Supp Table 8) targeting the plasmid backbone. The Mastermix was prepared as detailed before, except the concentration of each primer was 0.12 µM, and the annealing temperature was set to 62 °C. To determine whether producing the fluorescent protein had a fitness cost on the strains, their growth on PaFr$_{-CA_6}$ spent media was assessed. Precultures of the tagged and non-tagged strains were prepared in R2 broth as usual, inoculated to a final OD of 0.02 in 180 µL of spent media and incubated for 24 hours at 220 rpm and 28 °C. To check if either tagged strain had a fitness advantage over the other one, they were combined in different ratios, considering their OD, and grown in R2 broth under the same conditions as before. CFU's were counted after 24 h under UV light to differentiate each strain.

Infiltration and recovery of apoplast fluid was from *Flaveria* sp. leaves

For *F. linearis*, on two occasions four months apart (March 2020 and July 2020), five plants were grown and ten samples were collected (2 samples per plant). For *F. robusta*, three different sets of cuttings were sampled: one on March 2020 (4 different plants, sampled twice each), June 2020 (two different plants sampled twice) and April 2021 (2 different plants, sampled four and two times). For *F. trinervia*, four plants were sampled in March 2020, along with the other two species (2 samples per plant).

For the infiltration, the leaves were placed in a 60-cc syringe, which was filled with sodium phosphate (100 mM, pH 6.5). The plunger was pushed until the 50-cc mark to eject air, then pulled until the 55-cc mark and released back to the 50-cc mark; this was repeated several times until the leaves lost buoyancy. To recover the apoplast fluid wash (AFW), the leaves were placed on a sheet of parafilm, rolled around a 15-mL tube, and placed inside a 50 mL tube. After three minutes of centrifugation at 2500 x g, the recovered AFW was transferred to a clean 1.5 mL tube. After storage at -20 °C, the samples were spiked with an internal standard of deuterated amino acids and subjected to metabolomic profiling via untargeted UHPLC-HRMS (Supp Table 9).

Ultra-high performance liquid chromatography - high resolution mass spectrometry

Both the apoplast fluid wash from leaves (AFW) and the samples from the cross-feeding experiments were analyzed by untargeted metabolomics. Before injection, the samples were mixed with a standard mix of deuterated amino acids (U-D 98% Cell Free Amino Acid Mix from Cambridge Isotope Laboratories, Inc.) to a final concentration of 20 mg mix / µL sample (individual concentrations are shown in Supp Table 8). Ultra-high performance liquid chromatography coupled with high resolution mass spectrometry was carried out using a THERMO (Bremen, Germany) UltiMate HPG-3400 RS binary pump, WPS-3000 auto sampler which was set to 10 °C and which was equipped with a 25 µL injection syringe and a 100 µL sample loop. The column was kept at 25 °C within the column compartment TCC-3200. Chromatography column was used THERMO Accucore® C-18 RP (100 × 2.1 mm; 2.6 µm) using the following gradient of Eluent A (water with 2% acetonitrile and 0.1% formic acid) and Eluent B (pure acetonitrile), respectively at a constant flow rate of 0.4 mL/min: 0 min (100% and 0%), 0.2 min (100% and 0%), 8 min (0% and 100%), 11 min (0% and 100%), 11.1 min (100% and 0%), 12 min (100% and 0%). Mass spectrums were recorded with THERMO QExactive plus orbitrap mass spectrometer coupled to a heated electrospray source (HESI). Column flow was switched at 0.5 min from waste to the MS and at 11.5 min again back to the waste, to prevent source contamination. For monitoring two full scan modes were selected with the following parameters. Polarity: positive; scan range: 80 to 1200 m/z; resolution: 70,000; AGC target: 3 × 106; maximum IT: 200 ms. General settings: sheath gas flow rate: 60; auxiliary gas flow rate 20; sweep gas flow rate: 5; spray voltage: 3.0 kV; capillary temperature: 360 °C; S-lens RF level: 50; auxiliary gas heater temperature: 400 °C; acquisition time frame: 0.5 - 11.5 min. For negative mode, all values were kept instead of the spray voltage which was set to 3.3 kV.

Metabolomics data analysis

The raw data files were converted into mzML format in MSConvert (ProteoWizard, Version 3.0.19246-075ea16f5). To reduce the size of the files, we filtered them by peak picking with the Vendor algorithm, followed by threshold peak filtering using absolute intensity at 1.0E2. Next, we imported them into MzMine (version 2.40.1) for processing. The spectrums were first separated into positive and negative ionization modes. The baseline was corrected at 1.0 m/z bin width and using the asymmetric correction method. The peaks were then selected by centroid detection and by setting the noise level at 1.0E2. The chromatograms were built with the ADAP Chromatogram builder. The settings to build the chromatograms were slightly modified for each run by verifying the right integration of the deuterated amino acids peaks. The minimum group size in a number of scans was set to 2-3, the group intensity threshold was either 1.0E4 or 6.0E4, the minimum highest intensity was set at the same value as the group intensity, and the m/z tolerance was set at 0.001 m/z. The generated chromatograms were deconvoluted with the Wavelets (ADAP) algorithm, setting a signal to noise threshold of 10.0, a minimum feature height of 1.0E4, a coefficient/area threshold of 50, peak duration range 0-1 and a RT wavelet range of 0-0.1. Furthermore, the peaks were aligned together using the Join Aligner with m/z tolerance of 0.001, retention time tolerance of 0.1 min, and setting the weight for m/z and RT at 75 ppm and 25 ppm respectively. Finally, the matrix of negative and positive mode peaks, with their corresponding ID, m/z, retention time, and area were exported into R (version 3.6.0). Using an in-house script, uncommon (in less than 50% of the samples in the run) and small peaks (area <104) were filtered out. In the case of the AFW samples, the areas were corrected by the infiltration ratio.

To get a first overview of the distribution of the peaks across plant species, we conducted a principal coordinate analysis, constrained by plant species using Euclidean distance on the corrected peak matrix with the R packages phyloseq and vegan. Before plotting the data, the areas under the curve of each peak were transformed by glog2(x)= log2*{[x+sqrt(x2+a2)]/2}, where “a” is a constant with a default value of 1. We used this transformation as it has been shown to emphasize biological variation over technical variation in metabolomic data (Parsons et al., 2007). To compare the amino acids, the peak matrix was first annotated against a custom library of 570 compounds including all proteinogenic amino acids (MSMLS, Sigma Aldrich) which had been developed in the same equipment and UHPLC-MS method we used. This was carried out in R with an in-house script by setting the m/z tolerance at 0.002 and the RT tolerance at 0.2 min. The area of each amino acid was corrected towards its deuterated internal standard as follows:

$$\text{Corrected area} = \text{Area of amino acid in sample} * \frac{\bar{x}\ \text{area of deuterated amino acid in all samples in the run}}{\text{area of deuterated amino acid in sample}}$$

Next, we calculated the concentration of each amino acid, based on the known concentrations of the standard (Supp Table 8) and compared the amino acids concentrations in a heat map with the R package ComplexHeatmaps, using Euclidean distance and Ward’s algorithm to cluster the samples.

To find out which compounds had been taken up via cross-feeding, the spent media was analyzed before and after growth of the consumer isolates. To focus exclusively on uptake of large peaks, only peaks present in high amounts in the spent media (before growth of the consumers) were considered. This cutoff was defined for each dataset and ranged between 1.0E4 and 1.0E5. The log2 fold change, p-value and FDR were calculated for each peak. Metabolites with a fold change lower than -2 (after growth / before growth) and an FDR <0.05 were defined as significantly taken up. Venn Diagrams were created using the online tool Venny (version 2.1.0) and heatmaps were built in R with the package ComplexHeatmaps. The resulting peak tables were annotated against the custom library as described above.
